# A Complex Containing the Na^+^/K^+^ ATPase Regulates Epileptogenesis in *Drosophila melanogaster*

**DOI:** 10.1101/2023.09.22.559031

**Authors:** Derek M. Dean, Sarah Bermudez, Victoria Alkin, Ziming Wang, Huy Pham, Emily Kondo, Andrew Amini, David L. Deitcher

## Abstract

Epileptogenesis, the process through which the brain becomes seizure-prone, is not well understood. Previous work identified a novel gene in *Drosophila*, *julius seizure* (*jus*), that when mutated or developmentally knocked down, leads to epilepsy in adult *Drosophila*, providing a useful model for dissecting epileptogenesis. Here we report that when a GFP-tagged Jus was used as bait in a co-immunoprecipitation (co-IP) assay, a complex of 23 associated proteins was identified that included ATPalpha and Nervana 3, two subunits of the Na^+^/K^+^ ATPase. RNAi-mediated knockdown of ATPalpha, Nervana 3, or any of 8 additional complex proteins enhanced seizure susceptibility of *jus* mutants. The critical period for Jus expression to protect against epileptogenesis was further defined, occurring between pupal stages P4-7; remarkably, the *jus* seizure phenotype could be rescued by increasing neural activity of *jus*-expressing neurons during these mid-pupal stages, suggesting that altered neural activity in these neurons may contribute to the seizure phenotype. Our data support a model that, in wild type flies, a protein complex containing Na^+^/K^+^ ATPase and Jus prevents epileptogenesis, possibly by regulating neural activity.

## Introduction

Epilepsy is a neurological disorder characterized by unprovoked and recurring seizures that affects 1 in 26 people in their lifetime (Hesdorffer et al., 2011). Currently available anti-epileptic drugs (AEDs) can lessen symptoms in approximately 2 out of 3 patients. However, no effective treatment exists to prevent epileptogenesis, nor is the process well-understood beyond some preclinical models (Löscher, 2020; Löscher and Klein, 2021).

In rodent models, acute treatment with convulsants to mimic epileptic symptoms, electrical kindling to trigger epileptic foci, or genetic knockins of human disease variants have all provided valuable information about seizure mechanisms (reviewed in Marshall et al., 2021). However, modeling epileptogenesis in a rodent system has been difficult owing to the complexity of the mammalian brain, limited accessibility during embryonic development, and somewhat limited ability to target gene expression precisely in time and space.

In contrast, *Drosophila* have an even more sophisticated array of genetic and transgenic tools along with a modest-sized nervous system of about 140,000 neurons (Dorkenwald et al., 2024). Epilepsy models in *Drosophila,* known as bang sensitive (BS) mutants, undergo seizures in response to mechanical electrical or temperature shocks, or less effectively, to strobe light exposure (Benzer, 1971; Ganetzky and Wu, 1982; Lee and Wu, 2002; Pavlidis and Tanouye, 1995; Burg and Wu, 2012; Giachello and Baines, 2015; Dean et al., 2018). This behavioral output is easily scored, but there are challenges in using BS mutants to study epileptogenesis. Most BS mutations affect critical genes that are very widely expressed and affect neuronal function in most, if not all, cells (Royden et al., 1987; Pavlidis et al., 1994; Schubiger et al., 1994; Fergestad et al., 2006). Normally in *Drosophila*, suppressor/enhancer screens would be an effective way to identify genes that interact in a molecular pathway. Seizures, however, represent output from the entire CNS, and so genes may interact with the BS gene indirectly, enhancing or suppressing seizures by acting elsewhere in the CNS and/or at a different developmental period. This in mind, cell- and developmental stage-specific knockdown of interacting genes would more strongly support their role in an epileptogenic pathway.

*julius seizure* (*jus*), a recently identified BS gene, may provide a foothold to study epileptogenesis. *jus* encodes a predicted two-transmembrane protein. During pupal life, *jus* is expressed in selected neurons in the central brain, including the protocerebrum, and in a wide but specific pattern in the thoracic abdominal ganglion. In adults, Jus expression continues in the thoracic abdominal ganglion and central brain, with additional expression detected in the optic lobe and photoreceptors. Expression of Jus during pupal life is crucial to prevent epilepsy in the adult (Horne et al., 2017; Dean et al., 2018; Dean et al., 2026). In short, Jus has features that could make it valuable for studying epileptogenesis: namely, it has a critical developmental period for expression, and it is expressed in specific subpopulations of neurons.

The mechanism of Jus function is unclear. Other than its transmembrane spans, it has no domain that strongly predicts its function. Moreover, Jus has no known vertebrate homologue (Hu et al., 2011; Horne et al., 2017). Rather than conducting genetic interaction screens, we sought to identify evolutionarily conserved proteins that physically associate with Jus. We performed a large-scale co-immunoprecipitation with an epitope-tagged version of Jus, Jus-TGVBF, followed by SDS-PAGE and mass-spectrometry. From this analysis, 16 proteins were specifically co-immunoprecipitated with the tagged version of Jus and another 7 were found to be highly enriched in the animals expressing Jus-TGVBF as compared to controls. Using a *jus*-specific GAL4 to drive RNAi to each of the co-IP hits in a sensitized genetic background resulted in the identification of 10 genes with a role in epileptogenesis, all of which have orthologs in humans. The physical association of Jus with these highly conserved protein partners, the effect of knockdown of these protein partners on bang sensitivity in specific neurons, the developmental requirement for *jus* expression, and the interaction between neural activity and seizure sensitivity suggest an epileptogenic pathway in *Drosophila* that may inform studies of vertebrate epileptogenesis.

## Results

### Co-IP of Jus reveals protein partners

To identify specific protein partners of Jus, a large-scale Co-IP was performed. To express the bait, we used a transgenic fosmid line containing the *jus* locus tagged with GFP, Flag, and V5, which we abbreviate as *Fos{jus-TGVBF}*. This fusion construct has been shown to express functional Jus protein capable of at least partially rescuing the BS mutant phenotype, suggesting that its expression pattern overlaps that of endogenous Jus protein (Horne et al., 2017). Head extracts from the fosmid line (*w^1118^*; *Fos{jus-TGVBF}*) and from controls (*w^1118^*) were incubated with anti-GFP nanobody magnetic beads, purified, and eluted (Fig. 1). To confirm that the co-immunoprecipitation was able to recover Jus-TGVB from the lysate, a western blot was probed with anti-Flag antibody. A band of the predicted size of the tagged version of Jus was detected in the *w^1118^*; *Fos{jus-TGVBF}* lane but not in *w^1118^* lane (Figure 1). The remaining portion of the sample was separated by SDS-PAGE, digested within the gel, and subjected to mass spectrometry (Figure 1). Table 1 lists proteins that were unique to, or highly enriched in, the *w^1118^*; *Fos{jus-TGVBF}* extract relative to the *w^1118^* extract. As expected, Jus-TGVBF as recovered in the Co-IP experiment along with 16 other proteins that were only found in the Jus epitope-tagged animals but not in controls. An additional 7 proteins were highly enriched in the fosmid line as compared to the control (indicated in Table 1 with an asterisk). The most abundant pulldown specific to the *w^1118^*; *Fos{jus-TGVBF}* extract was the protein encoded by the rare exon 6a isoform of the alpha subunit of the Na+/K+ ATPase (Palladino et al., 2003). The beta subunit of the Na^+^/K^+^ ATPase, Nervana3, was also pulled down.

**Figure 1.**
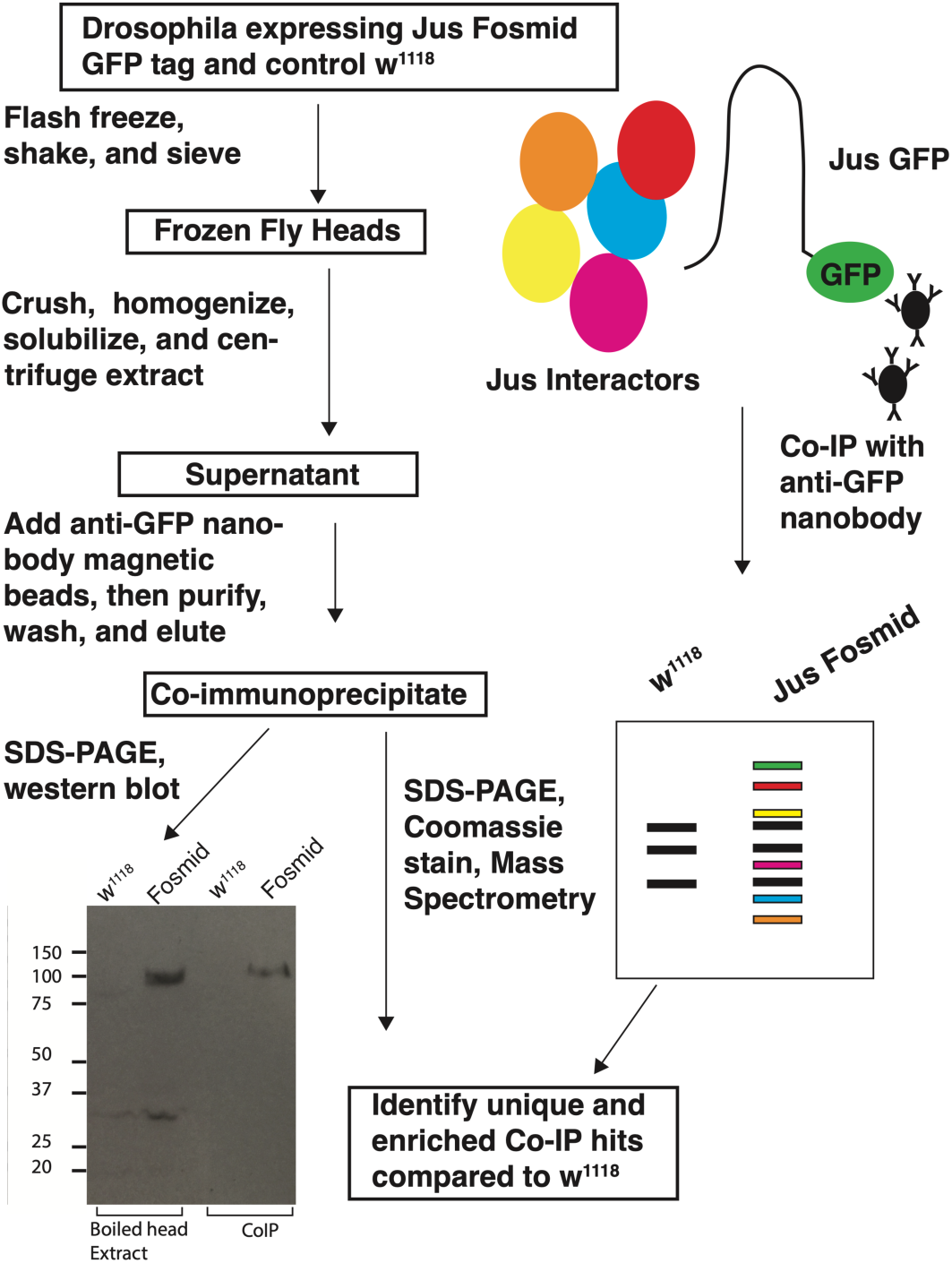
Identifying Jus protein interactors. Head extracts from *jus-TGVBF* (tagged with GFP, FLAG, and V5) and *w^1118^* control flies were used in a co-immunoprecipitation with anti-GFP nanobody magnetic beads. Purified extracts were run on SDS-PAGE gels; one gel was subjected to western blotting and probed with anti-Flag antibody, while other was stained with Coomassie, then co-immunoprecipitated proteins were identified with mass spectroscopy. Tagged Jus protein was detected on a western blot in boiled head extracts from *jus-TGVBF* flies and from the *jus-TGVBF* co-IP but not in *w^1118^*controls

**Table 1.** Co-immunoprecipitation hits sorted descending Sequest HT scores. Asterisked proteins were enriched relative to controls but not unique to tagged Jus sample.

| Proteins Recovered (Uniprot) | Fly Gene | Domains/function |
| --- | --- | --- |
| E1JIR4 | *ATPalpha*, exon 6A isoform | Na<sup>+</sup>/K<sup>+</sup> ATPase, alpha subunit |
| P13607* | *ATPalpha*, exon 6B isoform | Na<sup>+</sup>/K<sup>+</sup> ATPase, alpha subunit |
| A0A0B4KER6 | *Zasp52* | PDZ domain |
| P61857 | *betaTub85D* | beta tubulin |
| Q8MQS2 | *julius seizure*, isoform B | two-transmembrane domains |
| A0A0B4KHA7 | *julius seizure*, isoform D | two-transmembrane domains |
| Q0E9E2 | *Pyruvate carboxylase* | conversion of pyruvate to oxaloacetate |
| A0A0B4K7L1 | *Phosphofructokinase* | conversion of fructose-6-phosphate to fructose-1,6-bisphosphate |
| Q95U38 | *skpA associated protein* | subunit involved in succinate to succinyl-CoA |
| P02572 | *Act42A* | beta actin |
| E1JIC5 | *CG14995* | possible cytoskeletal protein |
| A8DYJ2 | *Hu li tai shao*, isoform P | adducin homolog |
| Q0E8G6* | *CG3689* | mRNA processing |
| M9PF61 | *Aldo-keto reductase 1B* | aldol reductase |
| A1ZAA5* | *Glutamate oxaloacetate transaminase 1* | involved in glutamate biosynthesis |
| Q9VH09 | *CG3999* | Glycine dehydrogenase |
| Q7K0L2 | *CG17754* | Ubiquitination, Kelch domain |
| M9PDX8* | *nervana3* | beta subunit of Na<sup>+</sup>/K<sup>+</sup> ATPase |
| Q9VFQ9 | *Dipeptidase B* | peptidase |
| A0A0B4KEY4* | *Patronin* | microtubule minus-end binding protein |
| Q8MZI3 | *CG10077* | RNA helicase, DEAD-box domain |
| P15357 | *RpS27A* | component of 40S ribosomal subunit, ubiquitin domain |
| P29843 | *Heat shock protein cognate 1* | misfolded protein binding activity |
| Q08473* | *squid* | hnRNPA family of RNA binding proteins |
| Q9VZ64* | *cg17333* | 6-phosphogluconolactonase |
| P48588* | *rps25* | component of 40S ribosomal subunit |

### Identifying A More Cell-Specific GAL4

To test the role of the Co-IP hits in bang-sensitivity, we opted to knockdown their expression in Jus-expressing neurons. Previous work demonstrated that using the 55G02-GAL4 line to drive expression of a wildtype *jus* transgene could effectively rescue the bang sensitive phenotype of a mutant *jus* allele (Horne et al., 2018). However, this rescue was not complete, and later it was found that overlap of 55G02-GAL4 expression with endogenous Jus expression was limited in the adult CNS (Dean et al., 2018). These findings suggested that 55G02 may not be the most effective enhancer to manipulate gene expression in the Jus-expressing neurons that affect bang-sensitivity. This in mind, we tested 55G02 alongside a series of additional *jus*-GAL4 lines (Figure 2A): First, all four GAL4 lines were tested for their ability to cause bang sensitivity by driving expression of a *jus*-RNAi construct, then these drivers were compared for their ability to rescue the *jus^iso7.8^* phenotype by expression of a wildtype *jus^+^* transgene. 90B09-GAL4, a construct containing the *jus* promoter region (Figure 2A), was the most effective driver for both experiments, causing 100% bang-sensitivity when driving *jus*-RNAi (Figure 2B), and fully suppressing bang-sensitivity when expressing UAS-*jus* in *jus^iso7.8^* mutants (Figure 2C), significantly better than the 55G02-GAL4 driver. In a separate study, we have confirmed that 55G02-GAL4 expression partially overlaps that of the Jus-GFP protein trap reporter Jus^GFSTF^ during pupal life (Dean et al., 2026).

**Figure 2.**
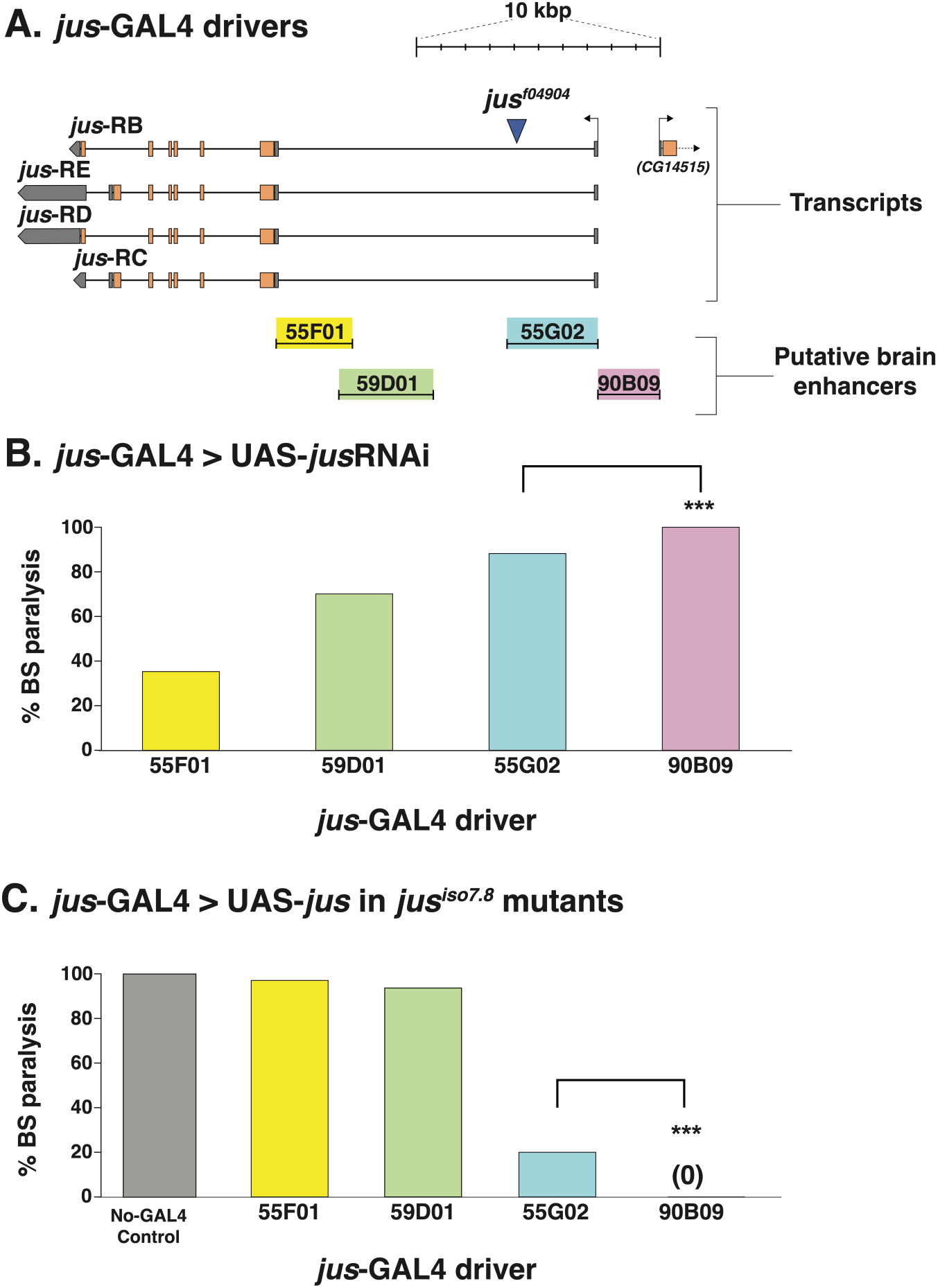
Identifying the most effective *jus*-GAL4 driver for *jus-*RNAi and for genetic rescue of *jus*. **(A)** *jus* locus and locations of the *jus*-GAL4 driver sequences we tested. Image was made with Adobe Illustrator using a rendering by GBrowse as a guide (Stein, 2013). Top right, 10 kilobase pair (kb) scale bar with 1 kb increments. Immediately below scale bar are the relative genomic positions of four known *jus* transcripts (*jus*-RB, -RC, -RD, and -RE) to one another and the 5’ end of the next gene to the right (*CG14515*). As in GBrowse, exons are indicated by rectangles, gray rectangles show 5’ and 3’-UTRs, orange rectangles indicate coding sequence, and introns are the lines between the exons. The promoter and enhancer sequences used to drive the four different *jus*-GAL4 constructs are indicated below the transcripts (see Materials and Methods for stock numbers). Driver sequences are color-coded to match them with the corresponding bars in the graphs below. **(B)** Bang-sensitivity of *jus*-GAL4 > UAS-*jus-*RNAi flies. *jus*-GAL4 drivers were used to express UAS-*jus-*RNAi in a *jus^+^*background. Of the four *jus*-GAL4 drivers, 55G02 and 90B09 induced the highest % bang-sensitivity (88% and 100% respectively). N>155 for 55G02 and 90B09-GAL4’s, *** p<0.0005, two-tailed Fisher’s exact probability est. **(C)** Bang-sensitivity of *jus* mutants expressing *jus*-GAL4 > UAS-*jus*. *Jus*-GAL4 drivers were used to express UAS-*jus* in *jus^iso7.8^*/*jus^iso7.8^* flies (“Homozygotes”). Of the four *jus*-GAL4 drivers, only 90B09 significantly completely suppressed the bang-sensitivity of *jus^iso7.8^* homozygotes. N>50 for 55G02 and 90B09-GAL4’s, *** p<0.0005, two-tailed Fisher’s exact probability test.

### RNAi of Protein Hits

To assess the role of the Co-IP proteins in bang-sensitivity, we drove expression of RNAi for each of the candidates using the *jus*-specific 90B09-GAL4 in a sensitized genetic background of *jus^iso7.8^/+* heterozygous animals. About 10% of *jus^iso7.8^/+* controls seized without knockdown of other genes, and RNAi of 10 candidates significantly enhanced bang-sensitivity over control levels: knockdown of *ATPalpha* modified this phenotype most strongly, with 100% of progeny demonstrating bang sensitivity. Other knockdowns that strongly enhanced bang sensitivity included *nervana3*, *CG10007*, *GOT1*, *RPS25*, *Act42A*, *CG17754*, and *Pfk* (Figure 3A). Using the same GAL4 line to express the same set of RNAi constructs but in a wildtype background, only knockdown of *ATPalpha* could cause bang sensitivity on their own, resulting in 100% of the flies being seizure prone. Initial RNAi knockdown of *nrv3* caused bang sensitivity, but repeated attempts failed to reproduce this effect.

**Figure 3.**
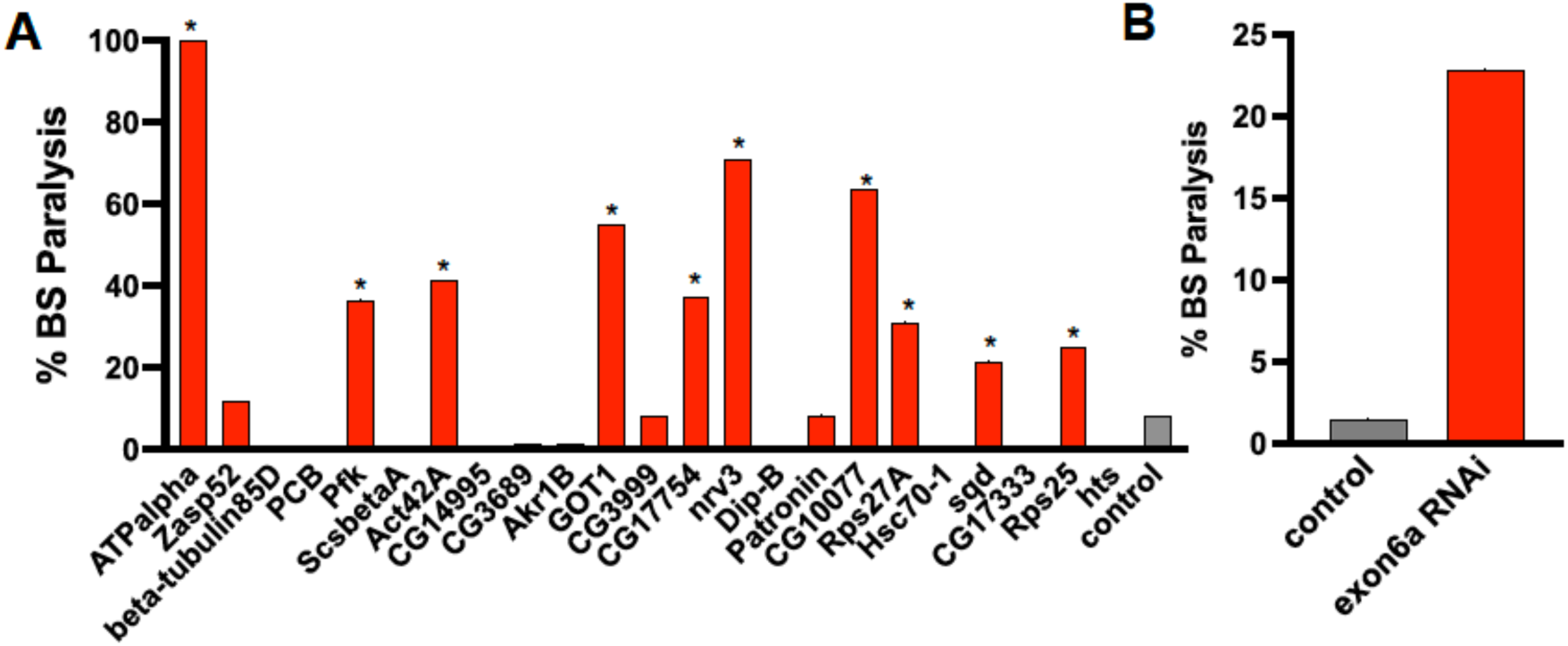
Bang-sensitivity enhancement by RNAi knockdown of Co-IP hits. **(A)** Bang sensitivity of *jus^iso7.8^*/*+* flies in which RNAi knockdown was performed on each of 23 Co-IP hits with the 90B09-GAL4 driver. RNAi knockdown of 10 different Co-IP hits significantly enhanced seizures. **(B)** Bang sensitivity of *jus^iso7.8^*/*+* in which 90B09>RNAi knockdown targeted exon6a of *ATPalpha*. N>20 for all genotypes, **** p<0.0001, *** p=0.0002, **p=0.0279, two-tailed Fisher’s exact probability test.

The *ATPalpha* RNAi construct used above targeted a 3’ UTR shared by many different *ATPalpha* transcripts, yet a rare isoform containing exon 6a encoded the most abundant pulldown from our Co-IP experiment. To determine if this particular isoform affects bang sensitivity, we generated a transgenic line to express RNAi specific to exon 6a. While exon 6a RNAi did not cause bang sensitivity in a *jus*^+^ background (not shown), it did significantly enhance bang-sensitivity in a *jus^iso7.8^/+* background, supporting its functional interaction with *jus* (Figure 3B).

### CG17754 affects Jus protein levels

How might knockdown of Jus protein partners enhance bang sensitivity? To begin to address this question, we selected *CG17754* for further investigation. This gene encodes a protein with BTB and Kelch domains. Such proteins often interact with Cullin3 to form E3 ligase complexes that mediate the ubiquitination of protein substrates (Shi et al., 2019), and so it could, directly or indirectly, regulate Jus expression and/or stability. Given our RNAi results, we first tested whether genetic mutants in *CG17754* could also enhance bang sensitivity.

P{EPgy2}*CG17754^EY05996^*, a transposon insertion in an intron of *CG17754*, roughly doubled the recovery time of *jus^iso7.8^*homozygotes (Figure 4A). We next used 90B09-GAL4 to drive expression of *CG17754-RNAi* in Jus-TGVBF-expressing animals. Head extracts from RNAi and control animals were separated by SDS-PAGE and subjected to western blotting. The blots were probed with an anti-FLAG antibody to detect the tagged Jus protein. Jus protein levels were significantly reduced in CG17754-RNAi animals, indicating that the presence of CG17754 might stabilize or prevent Jus degradation. Alternatively, it is possible that CG17754 normally increases Jus expression.

**Figure 4.**
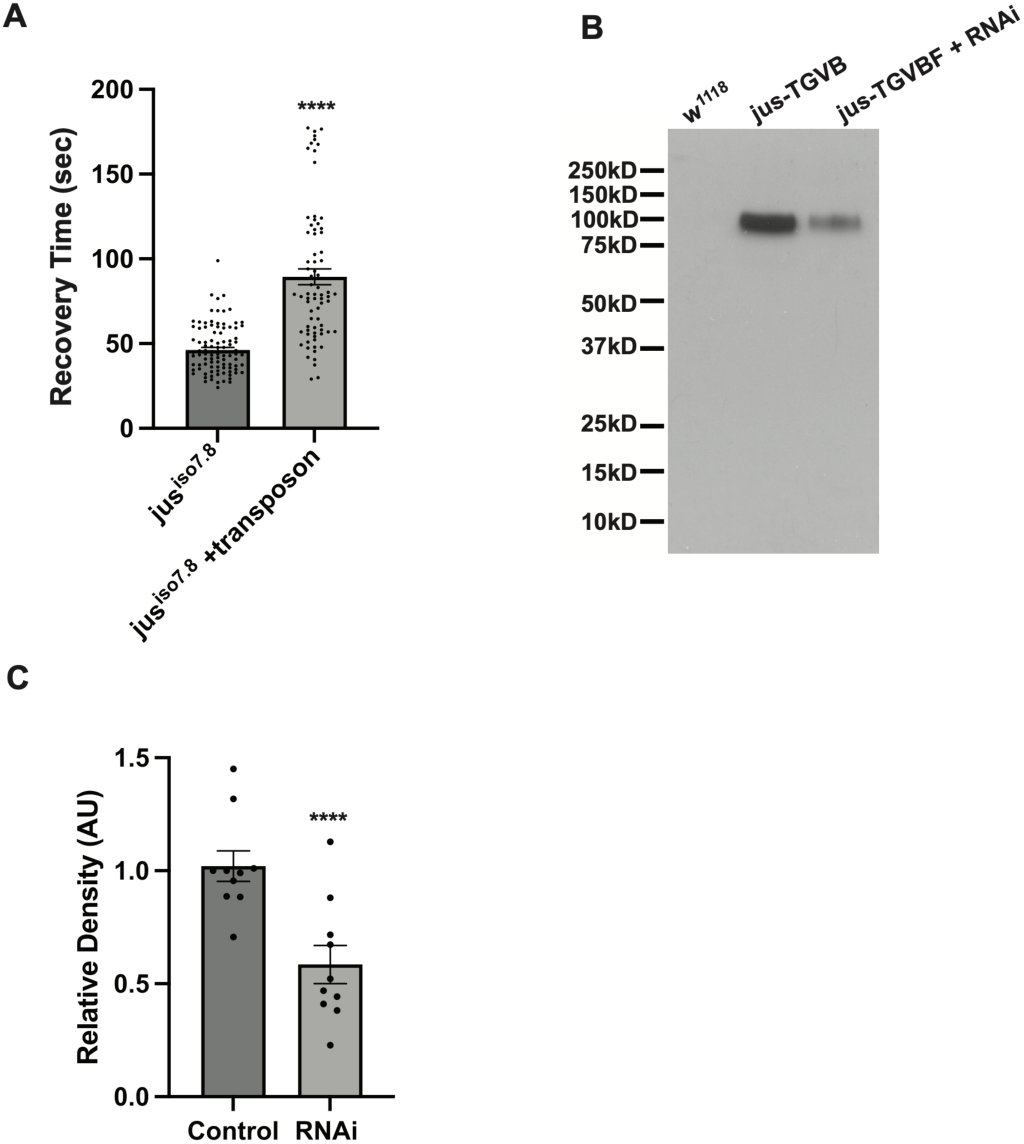
Reduced CG17754 levels results in enhanced seizures and a reduction of Jus protein levels. **(A)** The average seizure recovery times for the control flies homozygous for *jus*^iso7.8^ and flies homozygous for in *jus*^iso7.8^ and hemizygous for the *CG17754* transposon insertion P{EPgy2}*CG17754^EY05996^*. Dots represent individual data points. Error bars represent SEM. Control N = 99; transposon insertion and *jus*^iso7.8^ N = 74, **** p<0.0001. **(B)** Western blot comparing *Fos{jus-TGVBF}* flies to *Fos{jus-TGVBF}* with 90B09-GAL4>*CG17754-RNAi* knockdown. 10 fly heads were used in the sample for each lane. The *w^1118^* negative control lane did not produce bands with the anti-FLAG antibody. The bands produced by *Fos{jus-TGVBF}* with RNAi targeting *CG17754* are reduced relative to *Fos{jus-TGVBF}* alone **(C)** Comparison of relative densities of the bands from RNAi and Control. Knockdown of CG17754 resulted in a 41% reduction in Jus protein levels. N=10 for control and RNAi, **** p<0.0001, unpaired two-tailed t-test.

### Defining the Jus and ATPalpha critical phase

In a previous study of *jus*, animals expressing *Act5C*>RNAi-*jus* and GAL80^ts^ were incubated at 22°C then shifted to 29°C to induce RNAi expression at different developmental times. This experiment implicated a critical period for *jus* expression during the mid-pupal stage (Horne et al., 2016). However, the highly active *Act5C*-GAL4 construct was used to drive expression of *jus*-RNAi, and so GAL80^ts^ was unable to fully block bang-sensitivity, even if the animals were kept at 22°C throughout development. To improve the resolution of the assay, we repeated this experiment with one incubator set at 29°C as before but the other incubator set at 18°C instead of 22°C—this lower incubation temperature would be expected to decrease GAL4 efficacy while increasing the ability of GAL80^ts^ to block GAL4 activity (Rodriguez et al., 2012; Dean et al., 2016). Indeed, this updated protocol allowed more control over *jus*-RNAi expression (Figure 5): Control animals kept at 18°C throughout development had no bang-sensitivity, while animals kept at 29°C were fully bang-sensitive. Animals raised at 18°C then shifted to 29°C as late as pupal stage P4-5 were nearly as bang sensitive as animals raised at 29°C continuously. In contrast, a P6-7 18-29°C temperature shift resulted in bang-sensitivity in fewer than 30% of animals (Figure 5A), and even those that did seize recovered very quickly relative to animals that had been shifted earlier (Figure 5C). Further, shifting from 18°C to 29°C at P8-10 resulted in no bang sensitivity at all (Figure 5A). The reciprocal experiment of starting at 29°C and then shifting to 18°C revealed that shifts to 18°C prior to the wandering larval stage resulted in virtually no bang sensitivity. Shifting at wandering or P1-3 stages caused moderate bang-sensitivity, and nearly full bang sensitivity was reached if animals were shifted from 29°C to 18°C by P4-5 (Figure 5B). With the caveat that temperature shifts in either direction may require time to take effect, our results suggest that Jus expression between P4 and P7 is crucial for protecting against bang sensitivity.

**Figure 5.**
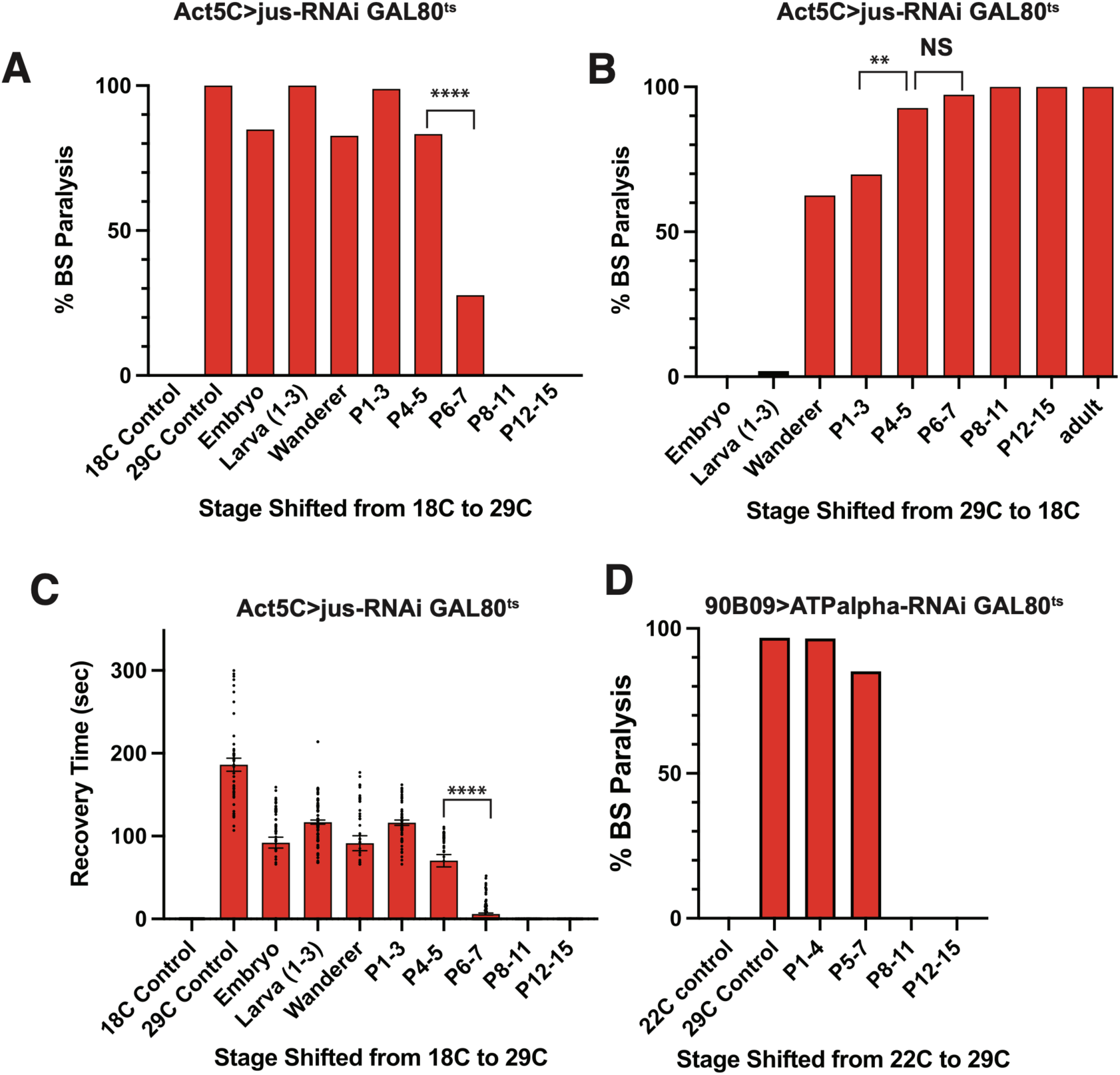
*jus* and *ATPalpha* have critical periods of expression between pupal stages P4-7 to protect against bang sensitivity. For **(A)** and **(B)** *Act5C>jus-RNAi* flies that also expressed *GAL80^ts^* were incubated at either 18°C or 29°C for the developmental times indicated. Flies kept at 18°C showed no bang sensitivity and flies kept at 29°C were all bang sensitive as predicted. **(A)** Animals switched from 18°C to 29°C as late as P4-5 exhibited 83% bang-sensitivity. However, animals switched at P6-7 were very significantly less bang sensitive, only 28% (p < 5X10^-12^, Fisher exact two-tailed probability test) and switching by P8 or later failed to cause bang-sensitivity. **(B)** Animals started at 29°C and switched to 18°C before the wandering larval stage showed virtually no bang sensitivity. Flies were significantly more bang sensitivity if they were shifted at P4-5 as compared to P1-3 (**p=0.0086, two-tailed Fisher’s exact probability test). But shifting at P6-7 was statistically no different (NS) from P4-5. **(C)** Recovery times from bang sensitive paralysis for part (A) sample groups. Shifting from 18°C to 29°C by P6-7 dramatically reduced recovery times, but as mentioned in A, animals shifted by P8 or later had no seizures at all. **(D)** 90B09>*ATPalpha-RNAi*, *GAL80^ts^* flies were incubated at 22°C until the indicated stage and then switched to 29°C. No flies seized if kept at 22°C for all of development, and all animals kept at 29°C seized. Nearly all animals seized when shifted to 29°C at P5-7 yet shifting by P8-10 or later failed to cause bang-sensitivity.

Having established a critical time window for *jus* expression, we wanted to determine if *ATPalpha*, the strongest Co-IP interactor, has a similar critical period. Using 90B09-GAL4 to drive *ATPalpha*-RNAi expression in the presence of GAL80^ts^, we raised animals at 22°C then shifted to 29°C at different pupal stages (Figure 5D). Animals raised at 22°C for all of development were not bang sensitive and those raised at 29°C continuously were 97% bang sensitive. Animals shifted from 22°C to 29°C at P1-4 were 96% bang sensitive, and those shifted to 29°C between P5-7 were still 85% bang sensitive. However, temperature shifts at P8 or later resulted in no bang sensitive adults. Thus, *ATPalpha* has a very similar critical period to *jus*.

### Increasing Activity During Pupal Stage Rescues Bang Sensitivity

As established earlier in this study, *jus* expression during the mid-pupal stage is critical to prevent seizures in the adult (Figure 5A-C). We hypothesized that a lack of Jus may alter the neuronal activity of *jus*-expressing neurons during this phase, leading to bang sensitivity. To increase neural activity, TrpA1, a temperature-sensitive, nonspecific cation channel (Pulver et al., 2009), was expressed using the 90B09-GAL4 line in a *jus^iso7.8^* mutant background. Animals were raised at 25°C then heated to 29°C at different stages of development to increase neural activity, and the resulting adult flies were tested for bang-sensitivity (Figure 6). Controls came out as expected: *jus^iso7.8^* 90B09>TrpA1 animals that were not heated remained highly bang-sensitive, and heating *jus^iso7.8^* mutants that carried UAS*-trpA1* but not a GAL4 driver at different developmental stages did not suppress their bang-sensitivity, except for some moderate suppression if animals were heated during the third instar stage (Figure 6A). However, heating *jus^iso7.8^* 90B09>TrpA1 P6-7 pupae until their eclosion rescued the bang-sensitivity of nearly 80% of the animals (Figure 6B), and the animals that did seize, recovered much more quickly (Figure 6C). In contrast, earlier or later heating had much less prominent effects. We also sought to reduce *jus*-expressing neuronal activity by expressing Kir2.1 or the light chain of tetanus toxin (Thum et al., 2006) with 90B09-GAL4, even at 18^0^C, but both treatments resulted in 100% lethality of the progeny.

**Figure 6.**
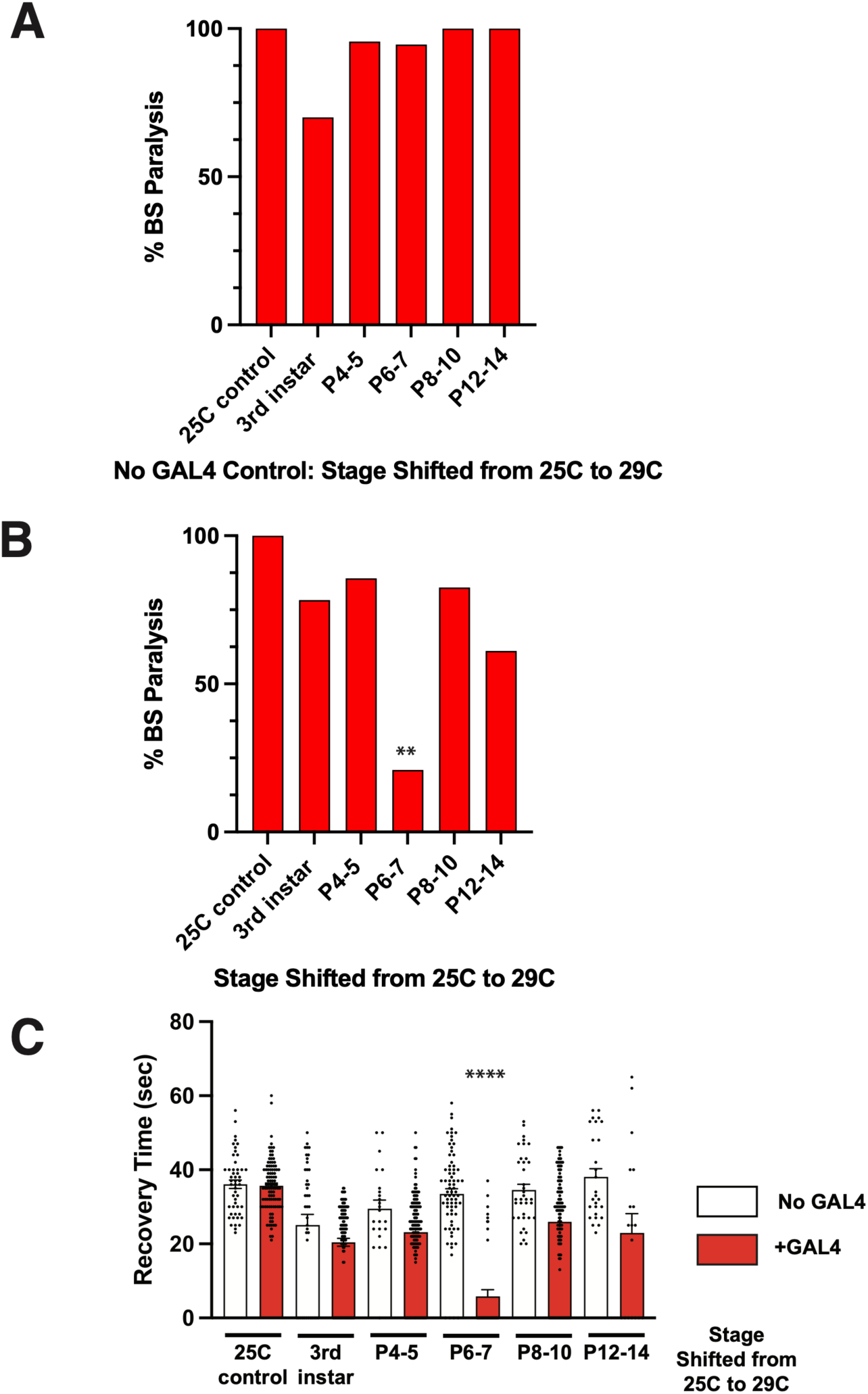
Developmental activation of TrpA1 rescued bang sensitivity. *TrpA1* was expressed in *jus*-expressing neurons in *jus*^iso7.8^ homozygotes (genotype: w; *UAS-TrpA1*/+; *90B09-gal4*, *jus*^iso7.8^ / *jus*^iso7.8^) and then shifted from 25^0^ C to 29^0^ C at different developmental times until eclosion to activate TrpA1. The adult flies were then tested for bang sensitivity and recovery times were recorded. **(A)** To control for temperature effects on bang sensitivity independent of TrpA1, *UAS-TrpA1*/+; *jus^iso7.8^ / jus^iso7.8^* (lacking GAL4) animals were shifted to 29°C at different developmental times. Heating third instars had a modest effect on bang sensitivity, but all the other time points were unaffected by the temperature shift. **(B)** Bang sensitivity is observed in all the *90B09>TrpA1; jus^iso7.8^ / jus^iso7.8^* flies raised at 25°C (25°C temperature control). Shifting to 29°C at P6-7 rescued 79% of the flies. In contrast, shifting at any other developmental stage did not cause anywhere near that level of rescue, two-tailed Fisher’s exact probability test, **p< 0.006 as compared to all other time points **(C)** Recovery times for the animals from part A and B. Although heating TrpA1-expressing animals at any developmental stage at least partially reduced recovery times, animals heated at the P6-7 stage had by far the fastest recovery, ****p<0.0001, one-way ANOVA with Tukey’s multiple comparison test.

## Discussion

This study identified 23 proteins that associate with Jus-TGVBF by co-immunoprecipitation (Table 1). Of the 23, 10 proteins were found to play a role in epileptogenesis by RNAi knockdown in *jus*-expressing neurons. The most abundant of the 23 proteins from the pulldown was the alpha subunit of the Na^+^/K^+^ pump, ATPalpha, and knockdown of this protein had the strongest effect on bang-sensitivity (Figure 3). Developmental knockdown of *jus* identified a critical stage of pupal stage P4-7 for epileptogenesis (Figure 5A-C), and knockdown of ATPalpha in *jus*-expressing neurons identified a very similar critical developmental stage (Figure 5D). Further exploration supported the idea that other proteins in the Jus complex are important for epileptogenesis. In particular, knockdown of CG17754, either directly or indirectly, reduced Jus protein levels, suggesting a mechanism by which CG17754 affects seizure sensitivity. Lastly, activation of TrpA1 in *jus*-expressing neurons demonstrates that the adult seizure phenotype can be ameliorated or rescued by affecting pupal neuron firing.

The main binding partner of Jus is the alpha subunit of the Na^+^/K^+^ ATPase. Mildly bang sensitive mutations in *Drosophila ATPalpha* have been previously reported (Schubiger et al., 1994; Sun et al., 2001; Palladino et al., 2003; Trotta et al., 2004; Ashmore et al., 2009). Our findings build on these studies in several ways: First, previous studies used genetic mutants, affecting the entire CNS and all other cells, while we specifically knocked down *ATPalpha* expression in *jus*-expressing neurons. Secondly, our results show a developmental effect of ATPalpha: Knockdown of *ATPalpha* after the P4-7 developmental window does not affect bang sensitivity. Lastly, the recovery time of *ATPalpha* genetic mutants ranges from 1-10 seconds (Schubiger et al., 1994; Ashmore et al., 2009), weaker than even the weakest of *jus* alleles. Mutations in the α subunit of the Na,K-ATPase can also lead to epilepsy in humans. Mutations the α 2 and 3 subunits of the Na,K-ATPase can cause migraine disorders known as Familial Hemiplegic Migraine (FHM) and Alternating Hemiplegia of Childhood (AHC), respectively. Epilepsy occurs in about 15-30% of FHM patients and in 50% of AHC patients (Vetro et al., 2021).

What aspect of Na^+^/K^+^ ATPase function is important for the seizure phenotype observed in *jus* mutants? One potential activity of ATPalpha that could lead to epilepsy is its core function of maintaining the negative resting potential. If *jus* mutations lead to lower activity of the pump, then the *jus* expressing neurons may have a more depolarized potential leading to a higher probability of neuronal firing. The TrpA1 experiment would seem to argue against this idea as increasing activity can rescue the seizure phenotype (Figure 6). An alternative is the ability of the Na^+^/K^+^ ATPase to respond to repetitive action potentials with a sustained long-lasting hyperpolarization (Pulver and Griffith, 2010). This phenomenon known as sodium-dependent afterhyperpolarization or AHP was observed in *Drosophila* motoneurons. It can act as a calcium independent integrator of spike number, suggesting a role in reducing excitability. AHP (also known as ultraslow afterhyperpolarization or usAHP) is conserved; it was found to regulate the swimming behavior of frogs and fish and the central pattern generator (CPG) in neonatal mice (reviewed in Picton et al., 2017). More crucially for epilepsy, it is also found to operate in neocortical and hippocampal pyramidal neurons (Gulledge et al., 2013). The circadian regulation of *ATPalpha* expression may also relate to bang sensitivity. Two peaks of *ATPalpha* expression are observed in the *Drosophila* visual system (Damulewicz et al., 2013), a region where Jus protein is also found (Horne et al., 2017). Two peaks of bang sensitivity were also found in *jus* mutants, that correspond to peak expression levels of *ATPalpha* (Huertas-Radi et al., 2026).

A host of other proteins also associated with Jus, including CG17754. This protein has Kelch and BTB domains which often function in the ubiquitin degradation pathway (Shi et al., 2019). We investigated how CG17754 may enhance seizures and discovered that it either directly or indirectly regulates Jus protein levels (Figure 4B). Initially our expectation was that a protein involved in the ubiquitin pathway would promote Jus degradation rather than prevent it. It may be that CG17754 binds Jus protein but does not transfer it to the degradation pathway unless it is appropriately signaled. Alternatively, CG17754 may negatively regulate another protein that normally degrades Jus, and, in its absence, the other protein is free to degrade Jus. Given the genetic interaction of *CG17754* with *jus*, and the striking effect *CG17754* has on Julius Seizure protein levels (Figure 4). The physical and functional interactions between Jus and at least 10 of its protein partners—especially ATPalpha, Nrv3, and CG17754—strongly support the specificity of the co-immunoprecipitated proteins and their importance in epileptogenesis. How the other Jus proteins influence seizures will be the subject of further study.

We also investigated whether changes neural activity during the pupal stage could affect the *jus*-dependent seizure phenotype. Increasing neural activity in *jus*-expressing neurons with the temperature dependent TrpA1 ion channel during pupal stages P6-7 was able to fully rescue the seizure phenotype of approximately 80% of flies (Figure 6). This could suggest that *jus* mutants have a reduced firing ability during the pupal stage leading to an adult seizure phenotype. However, the high firing rate induced by TrpA1 and the extended period of this treatment is unlikely to replicate the normal firing pattern of *jus*-expressing neurons. Nonetheless, this elevated activity did rescue the *jus^iso7.8^* bang sensitive phenotype in approximately 80% of the animals and reduced the seizure duration dramatically in those not fully rescued.

In summary, the identification of a protein complex that acts in specific neurons to affect epileptogenesis represents a substantial advance in our understanding of how the CNS regulates neuronal excitability to prevent epilepsy. Although Jus does not appear to be conserved in vertebrates, many of the proteins in its partner complex are conserved in humans and may represent targets for the development of new anti-epileptic drugs. Further investigations of each Jus protein partner, and of the pupal neurons that express the complex, should yield new insights into the still mysterious process of epileptogenesis.

## Acknowledgments

We thank Sheng Zhang for his advice on co-immunoprecipitation protocol design, Elizabeth Anderson for her assistance in sample preparation for mass spectroscopy, and Ievgen Motorykin for proteomics data analysis. We also thank Bruce Johnson for insightful comments on the manuscript. We thank Katie McBride for preliminary studies with jus and TrpA1 and Caressa Swartz for her work on Tetx-LC and Kir2.1. This work was in part supported by the Triad foundation and from a grant from College of Arts and Sciences, Cornell University.

## Author contributions

Conceptualization, D.L.D. and D.M.D.; Analysis and Investigation, S.B., D.M.D, V.A. Z.W., H.P, E.K., A.A., and D.L.D.; Writing, D.L.D. and D.M.D; Supervision, D.L.D. and D.M.D.

## Materials and Methods

### Fly Stocks

Fly stocks were maintained on a glucose, yeast, and cornmeal agar media at 22-25 °C in plastic vials unless otherwise noted. Most stocks were obtained from the Bloomington Drosophila Stock Center (BL) (NIH P40OD018537) or from the Vienna Drosophila RNAi Center (V); their stock numbers will be described later in this Materials and Methods. The original *w; jus^iso7.8^* stock was a gift from Mark Tanouye. When applicable, *w^1118^* (BL #5905) was used as a control. For neuronal activity perturbations, stocks *w; UAS-trpA1* (BL# 26263), *w; P{w[+mC]=UAS-TeTxLC.tnt}E2* (BL#28837), and *w; P{w[+mC]=UAS-Hsap\KCNJ2.EGFP}7* (BL#6595) were used in the genetic backgrounds described. Crosses were performed at 25 °C with 12-hour light-dark cycle unless otherwise stated.

### Generation of Transgenics

P{GMR90B09-GAL4}, a plasmid containing the *GAL4* gene fused to the *jus* promoter region, was kindly provided by Heather Dionne at the Janelia Research Campus (Pfeiffer et al., 2008). P{GMR90B09-GAL4} was sent to Genetivision (Houston, TX), which generated a new *jus*-GAL4 transgenic line by inserting the construct into a docking site at 68A4. Antisense oligos targeting exon 6a (shown in red) were designed using the SnapDragon (https://www.flyrnai.org/snapdragon). The exon 6A RNAi construct was made by first heating 10uM of Trip oligos CTAGCAGTAGATCGATAACCTAGTCAACGTAGTTATATTCAAGCATACGTTGACTAGGTTATCGATCTGCG and AATTCGCAGATCGATAACCTAGTCAACGTATGCTTGAATATAACTACGTTGACTAGGTTATCGATCTACTG in 10 mM Tris pH7.5, 0.1 M NaCl, 1 mM EDTA at 95 °C in a heat block then turning it off to cool. The resulting annealed primers were added to NheI/EcoRI-cut Valium 20 vector and ligated with T4 DNA ligase and transformed into NEB Express competent cells. Colonies were grown up and miniprep DNA was screened by PCR for inserts with primers ACCAGCAACCAAGTAAATCAAC and TAATCGTGTGTGATGCCTACC. Two minipreps were sequenced to confirm the sequence of the insertion. The resulting vector was used to inject *y[1] w[1118]; PBac{y[+]-attP-3B}VK00002* embryos and obtain an insertion at 28E7 (Bestgene, Chino Hills, CA).

### Co-immunoprecipitation/Mass Spectrometry

Flies from 20 bottles of *w^1118^*; Fos{jus-TGVBF} (Horne et al., 2017) and control *w^1118^* (approximately 40 ml of packed flies from each genotype) were flash frozen in liquid nitrogen. Flies were stored in 50 ml polypropylene tubes at −80 °C. The tubes were then rapidly removed from the freezer, shaken vigorously to shear off the heads, and poured into a cold 710-micron brass sieve (Humboldt Mfg, Co. Chicago, IL). Sieved heads were ground using a chilled mortar and pestle on dry ice. The resulting head powder was poured into a 15 ml Pyrex homogenizer containing 7 mL of 1x lysis buffer (20 mM Tris pH 7.5, 300 mM NaCl, 1 mM EDTA, 1% Triton X-100) that contained 1X protease inhibitor complete tablets, EDTA-free (Roche). Homogenate was incubated on ice 25 minutes, placed on a rotator at 4 °C for 30 minutes, spun at 16,000 g for 10 minutes, then the resulting supernatant was spun again at 16,000 g for 5 minutes. The resulting supernatant was then spun through a spin filtration column (Ultrafree-MC-HV, Millipore, Cork, IRL) at 12,000 g for 10 minutes. 250 μL of pre-rinsed anti-GFP nanobody magnetic beads slurry (Chromotek) in 10 mM Tris pH 7.5, 150 mM NaCl, 0.5 mM EDTA were added to each sample containing filtered homogenate. Tubes were incubated overnight at 4 °C on a rotator (Fisher hemocytology mixer). Beads were captured with a magnetic separation rack (New England Biolabs), washed twice with chilled buffer (10 mM Tris 7.5, 150 mM NaCl, 0.5 mM EDTA, 0.1% Triton), eluted with 50 μL of SDS loading buffer (Laemmli, 1970), and heated to 95 °C for 10 minutes. The resulting solution was spun through a spin column (Ultrafree-MC-HV, Millipore, Cork, IRL) at 12,000 g for 5 minutes, then stored at −20 °C. 15 μL each of the *w^1118^*; Fos{jus-TGVBF} and *w^1118^*samples were run on a 0.75 mm 10% SDS-PAGE gel with 10 μL molecular weight marker (BIO-RAD Precision Plus Protein Standards, Dual Color). After electrophoresis, the gel was blotted onto an Immobilon PVDF membrane and probed with anti-FLAG as described below in western blotting (see below) to confirm the immunoprecipitation of Jus. The remaining 35 μL of each sample was run on a 1.5 mm 10% SDS-PAGE gel with 10 μL molecular weight marker, stained with colloidal Coomassie Blue (BIO-RAD) overnight, and destained in deionized water for 3 hours. The co-IP products in this gel were excised, subjected to digestion in-gel with trypsin, and then subjected to nano LC-MS/MS 90-120 min gradient on an Orbitrap Fusion. Proteins were identified based upon at least two different peptides.

### RNAi

For RNAi knockdown of *jus* expression, the following *jus* enhancer GAL4 lines were used (described in Pfeiffer et al., 2008): 55F051 (BL# 39124), 55G02 (BL# 46070; no longer available from BDSC but see Horne et al, 2017 and Dean et al., 2018), 59D01 (BL# 46416) and 90B09 (BL# 607421). Each of these *jus*-GAL4 lines was crossed to the UAS-*jus*RNAi line *P{y[+t7.7] v[+t1.8]=TRiP.JF03192}attP2* (BL#28764; Ni et al. 2009). Progeny containing both transgenes were tested for bang sensitivity. For developmental *jus*RNAi studies, an *Act5C-GAL4* (BL#3954) construct was recombined onto the UAS-*jus*RNAi third chromosome described above, then balanced over TM6C *Sb Tb*. The resulting line was crossed to *w[*]; P{w[+mC]=tubP-GAL80^ts^; TM2/TM6B, Tb* (BL#7108), and the resulting progeny were incubated at either 18^0^C or 29^0^C. Controls were maintained at one temperature or the other throughout development, while experimental groups were shifted from one temperature to the other at a specific developmental stage. GAL80^ts^ suppresses GAL4 function at 18^0^C but is inactive at 29^0^C. Therefore, the UAS-*jus*RNAi construct is expected to be expressed 29^0^C but not at 18^0^C (McGuire et al., 2004; Dean et al., 2015). *P{w[+mC]=tubP-GAL80^ts^/+; Act5C-GAL4, P{y[+t7.7] v[+t1.8]=TRiP.JF03192}attP2*/*TM2 or TM6B, Tb, Hu* animals were selected then vortex tested as described below (**Vortex Testing**). At least 52 animals were tested for each time point. For the RNAi knockdown of *ATPalpha*, 90B09-GAL4 was crossed to *y[1] sc[*] v[1] sev[21]; P{y[+t7.7] v[+t1.8]=TRiP.HMS00703}attP2* (BL# 32913; Perkins et al., 2015). Resulting progeny were tested for bang sensitivity as described below (**Vortex Testing**). The RNAi lines for each of the co-IP hits are listed in Table 2.

**Table 2.**
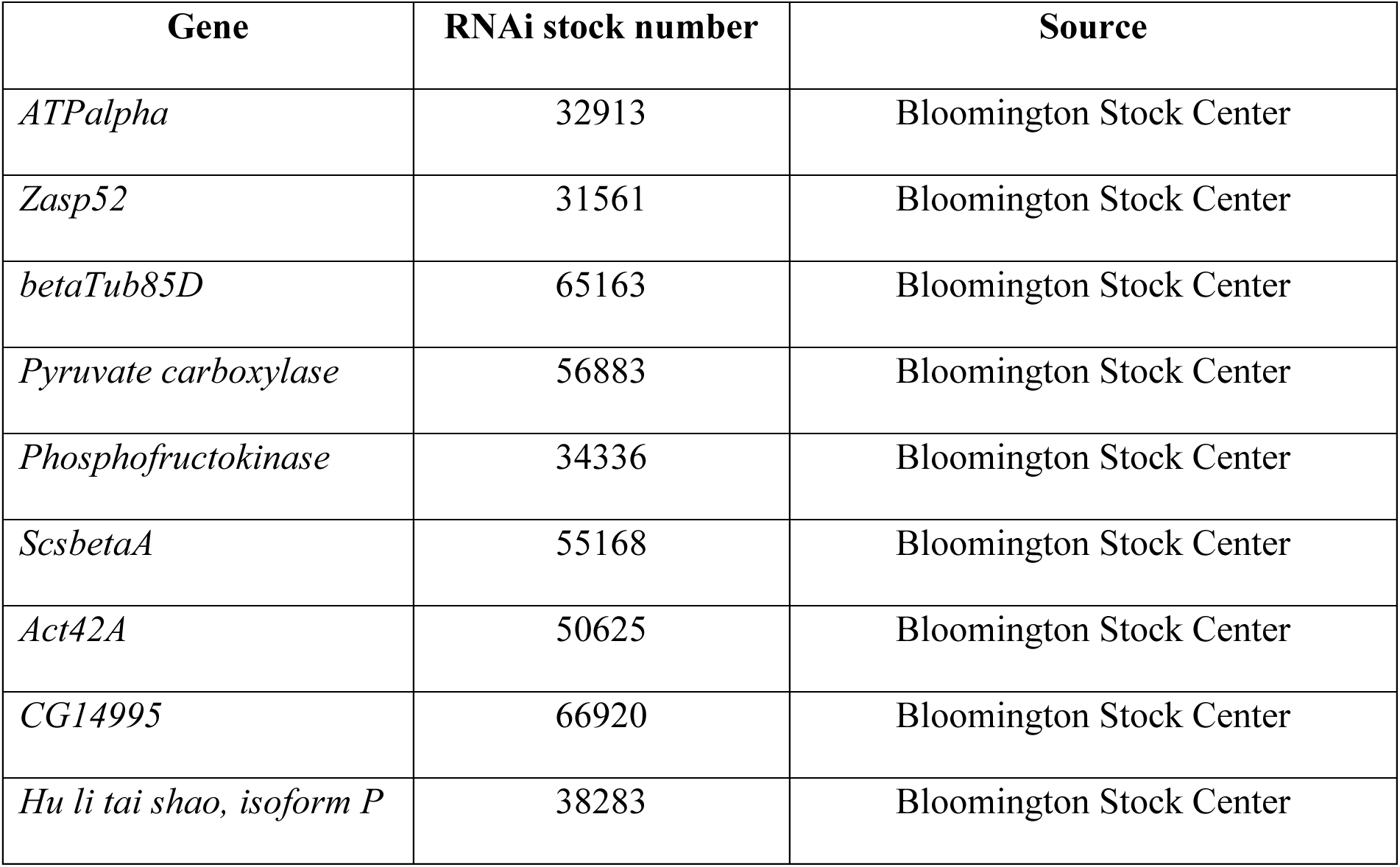
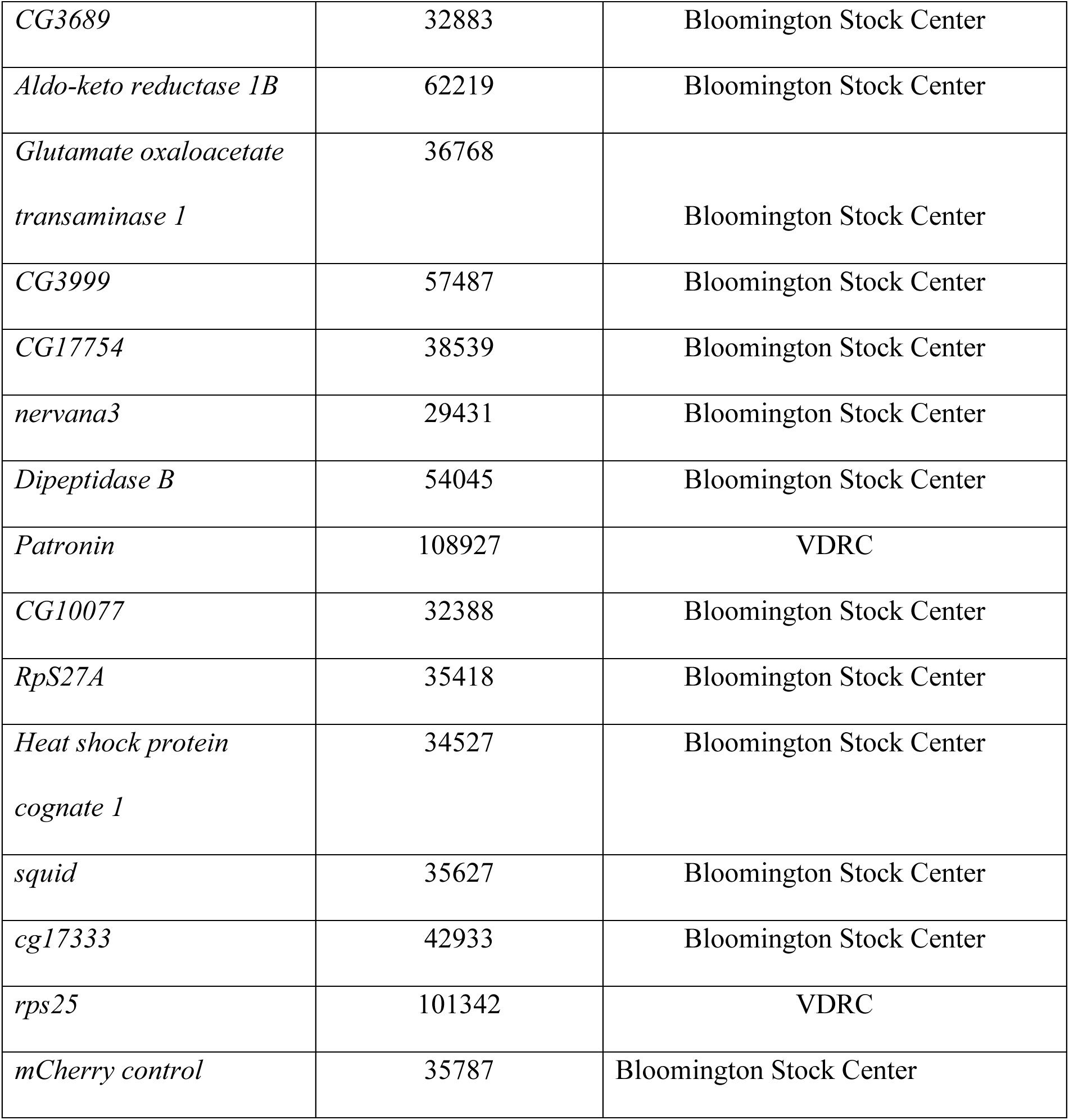
RNAi stocks used to test Co-IP hits.

### Genetic Rescue

A *UAS-jus* transgene insertion on the third chromosome was recombined with *jus^iso7.8^* as previously described (Horne et al., 2017). In separate crosses, 4 *jus*-GAL4 insertions described in the RNAi section—55F01, 55G02, 90B09, and 59D01 —were recombined onto the *jus^iso7.8^* chromosome. The UAS*-jus jus^iso7.^*^8^ stock was crossed to our 4 different *jus-*GAL4 *jus^iso7.8^* stocks. UAS*-jus jus^iso7.8^ /jus-*GAL4 *jus^iso7.8^* progeny were selected, allowed to recover from CO_2_ for 24 hours, then vortex tested as described below. A cross of the UAS*-jus jus^iso7.8^ to jus^iso7.8^ (*with no GAL4 driver) was used as a control.

### Western Blotting

To detect protein levels of Jus in *CG17754* RNAi knockdown animals, *Fos{jus-TGVBF}; 90B09-GAL4* animals were crossed to *CG17754-RNAi* (BL #38539, see Table 2) and the resulting *Fos{jus-TGVBF}/+; 90B09-GAL4/CG17754-RNAi* flies were selected. 10 heads of RNAi and control Fos{jus-TGVBF}/+; 90B09-GAL4/C+ were homogenized in SDS-PAGE loading buffer, separated on 10% polyacrylamide gels, then blotted on Immobilon P filters (Millipore). Filters were probed with anti-FLAG (M2, Sigma, St. Louis, MO), diluted 1:1000, then 1:10,000 donkey anti-mouse HRP (Jackson Immunoresearch) secondary antibody.

Detection was with either SuperSignal West Pico Plus or West Atto Chemiluminescent Substrate (Thermo Scientific), and imaging was with CL-XPosure Film. Developed film was digitally scanned and signals were quantitated with Fiji image analysis software (Schindelin et al., 2012). Means were determined from 3 western blots from 10 independent control and RNAi samples.

### Vortex Testing

For testing seizure behavior, adult flies were anesthetized with carbon dioxide and placed in vials containing food (up to roughly 25 flies per vial). After at least 24 hours after CO_2_ exposure, the flies were tested for bang-sensitivity by vortexing between ZT1-ZT4 in an inverted food vial for 10 seconds using a vortex at maximum speed. The number of flies that seized in response to the stimulus was recorded. Fly age was controlled between the sample groups of a given experiment: For experiments employing *jus-*RNAi and UAS-*jus*, adult flies were tested 1-3 days post-emergence, but for protein partner RNAi experiments, adults were tested at 3-7 days post-emergence.

### Statistics

Statistical analysis was performed with GraphPad Prism 10. Tests and their p-values are described in their associated figure captions.

